# Disentangling neural traveling waves from causal information flow

**DOI:** 10.1101/2025.02.25.640049

**Authors:** Andrea Alamia, Antoine Grimaldi, Frederic Chavane, Martin Vinck

## Abstract

In many behavioral conditions, neural activity manifests itself within and across brain regions as traveling waves, revealing the importance of analyzing spatiotemporal dynamics in electrophysiological data. Most methods detect traveling waves by measuring spatial phase gradients, i.e., monotonic and ordered phase changes through space. It is unclear, however, how these traveling waves relate to the causal directionality of information flow. Here, we analyze systems of coupled nodes with an external input to one node. We demonstrate that the phase ordering in traveling waves does not always correspond to the direction of effective information flow. We show that discrepancies can emerge in the case of systems with delays and inhibitory influences. As a methodological solution, we show that Granger causality analysis can, in linear systems, recover the directionality of the information flow. We propose a new measure called DIFF, the Directional Information Flow Field. DIFF is constructed by analyzing directed causal influences in space and time between neighbours, yielding a vector field. As a proof of principle, we show that, in a 2D field where a connected network is perturbed by an external input, the Divergence of DIFF can identify the spatial source of the perturbations. We propose that causal inference methods provide complementary information to phase-based traveling wave methods in analyzing system dynamics and information flow.

## Introduction

Recent experimental work demonstrates that neural activity propagates between and within brain regions as traveling waves at macroscopic and mesoscopic scales (Muller et al. (2014, 2018); Ermentrout and Kleinfeld (2001); Bhattacharya et al. (2022a); Rubino et al. (2006); Hatsopoulos et al. (1998); Troy and Shusterman (2007); Benucci et al. (2007); Richardson et al. (2005); Campbell et al. (2024); Pesaran et al. (2018)). Therefore, it is important to consider the spatial and temporal components in neural recordings to thoroughly characterize the putative role of such activity (Zhang et al. (2018); Lubenov and Siapas (2009); Patel et al. (2012); Davis et al. (2020); Zeng et al. (2024); Galas et al. (2025); Luo et al. (2021); Ye et al. (2023); Bhattacharya et al. (2022b); Reynaud et al. (2012); Chemla et al. (2019a)). Several methods have been proposed in the literature to identify traveling waves and characterize their properties (Das et al. (2023); Alexander et al. (2013); Alamia and VanRullen (2019); Schwenk and Alamia (2024); Davis et al. (2020); Muller et al. (2014)). Most of these methods rely on phase estimation to quantify the spatial gradients in a given frequency band and have provided relevant insights into the mechanisms associated with traveling waves.

Traveling waves are, in essence, spatial phase gradients, i.e., monotonic and ordered phase changes through space, and as such, they do not directly measure, and are not designed to extract, causal influences. It is therefore unclear how traveling waves relate to the directionality of effective information flow, and under which conditions these two measures provide different and complementary results. There are several methods that have been specifically designed for measuring the information flow based on causal influences, such as Granger Causality (GC) (Granger, 1969) (for capturing linear causal influences) and Transfer Entropy (Duan et al., 2013) (for capturing both linear and non-linear causal influences). However, it is unclear how an emergent pattern of phase relationships relates to these directional causal influences.

Here, we investigate the relationship between phase-based traveling waves and causal inference methods using different network configurations, in which we vary the sign and delay of the connections between nodes. Networks were composed of (linear) vector autoregressive (VAR) systems (Seth et al., 2015; Vinck et al., 2015; Pourahmadi and Noorbaloochi, 2016; Gorrostieta et al., 2013; Ghosh et al., 2019; Marinazzo et al., 2011; He et al., 2019; Dowdall and Vinck, 2023). All models were fed with a perturbation to the input node, which will generate information flow away from the input node and, as expected, should generate phase-based waves propagating in the same direction. In this way, we could compare propagating waves with some ground-truth flow of information. We used GC analysis to extract the causal direction of the information flow, which is supposed to extract the actual flow of information. This analysis was then compared with phase-based methods(Brovelli et al., 2004a; Bastos et al., 2015; Seth et al., 2015; Vinck et al., 2015; Ding et al., 2006). We show that discrepancies between phase-based and causal inference methods can emerge when the connections are partially inhibitory or contain asymmetric delays. We further show that GC analysis can, for the analyzed linear systems, recover the directionality of the information flow. We then propose a new measure called DIFF, the Directional Information Flow Field. DIFF is constructed by analyzing directed causal influences in space and time between neighbours, yielding a vector field. As proof of principle, we show that, in a 2D field where a connected network is perturbed by an external input, the Divergence of DIFF can accurately identify the spatial source of the perturbations.

The following sections of the paper are organized as follows: we first describe the mathematical formalism used to define the VAR models and the four connectivity conditions we investigated (i.e., from or to the external perturbation, either excitatory or inhibitory, see Figure 1A). We then define cases for a 1D VAR process with different symmetric and asymmetric temporal delays and one model inspired by a previously proposed predictive coding model (Alamia and VanRullen, 2019). In all cases, we consistently compare the results of the phase-based analysis with the GC analysis. We then extend these results from 1D to 2D cases, and develop and test the DIFF method to characterize the spatiotemporal organization of causal influences in different conditions.

**Figure 1:**
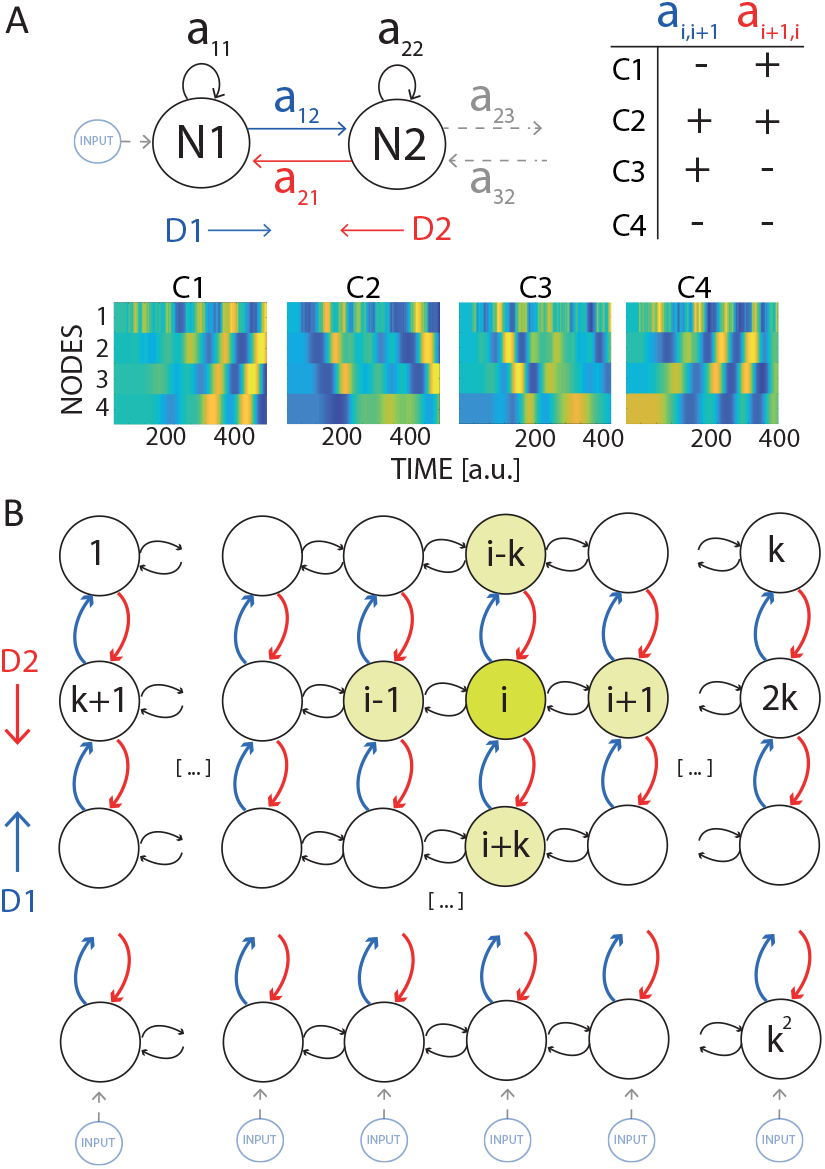
Figure 01 A) Schematic representation of the VAR models, in which subsequent nodes are connected via parameters *a*_*i,k*_. We considered four cases in which the connections in both directions have either positive or negative signs, as summarized in the table. We defined D1 as the direction from the perturbed node, and D2 as the opposite direction (in blue and red, respectively). The lower panels show 2D representations of the activity in the four cases of the Predictive Coding implementation with 4 nodes. The x-axis and the y-axis represent the time and the levels (i.e., nodes), respectively. The color (in arbitrary units) reflects the activity over time in each node. B) Schematic representation of the two-dimensional VAR model composed of *k*^2^ nodes. Each node is connected to its neighbors. A perturbation to the nodes in the lowest edge defines the direction D1 and D2 parallel to the perturbation (in red and blue in the figure). As in the 1D case, we defined four conditions by varying the signs of the D1 and D2 connections.

## Results

We first investigate 1D models where an external input was introduced in one node, generating a groundtruth information flow outward from this node, propagating through either excitatory (E) or inhibitory (I) connectivity (defining four conditions, as shown in Figure 1A). Further below, we also consider 2D models (Figure 1B). We will now systematically compare how GC and phase-based analyses capture the true direction of the information flow in different cases.

### Phase-based for the 1-D VAR(1) case

For the 1D case shown in Figure 1A, we first analyzed the phase differences between subsequent nodes. As the first node was perturbed by the external input, we expected traveling waves (TW) to emanate in Direction 1 from the first to the last node. However, as seen from the lower panels in Figure 1A, the direction of TWs was highly dependent on the sign of connectivity between the nodes. In particular, TWs appeared to emanate in Direction 1 from the first (lower) node to the last (higher) node when the connections in Direction 1 (i.e. *a*_*i,i*+1_ *>* 0) were excitatory. By contrast, TWs appeared to emanate in Direction 2 when the Direction 1 connections were inhibitory (i.e. *a*_*i,i*+1_ *<* 0).

To systematically quantify these differences, we computed the relative phase differences between subsequent nodes. We found a negative phase difference between all subsequent nodes when the *a*_*i,i*+1_ coefficient was negative (i.e., Cases 1 and 4), indicating a propagation backward from higher to lower nodes. We observed the opposite direction flow when the *a*_*i,i*+1_ coefficients (Direction 1) were positive (in the second and third cases, i.e., a positive phase difference between subsequent nodes. This result was supported by a Von Mises test (V-test) for non-uniformity of circular data considering the mean direction in any pair of nodes (in all conditions and pair of nodes, we obtained for k=3 all V-value *>* 250 and *p <* 0.0001; and for k=7 all V-value *>* 180 and *p <* 0.0001). We found consistent results for k=3 and k=7 nodes (Figure 2A,B) All in all, the results from the phase-based analysis suggest that the direction of propagation, as measured by the phase difference between subsequent nodes, depends on the sign of the *a*_*i,i*+1_ coefficient. A positive influence from the *i* to the *i* + 1 node determines the direction of phase-based propagation from lower to higher nodes (i.e., in direction D1, 1A), whereas a negative modulation produces a direction flowing in the opposite direction, from higher to lower regions (i.e., direction D2).

**Figure 2:**
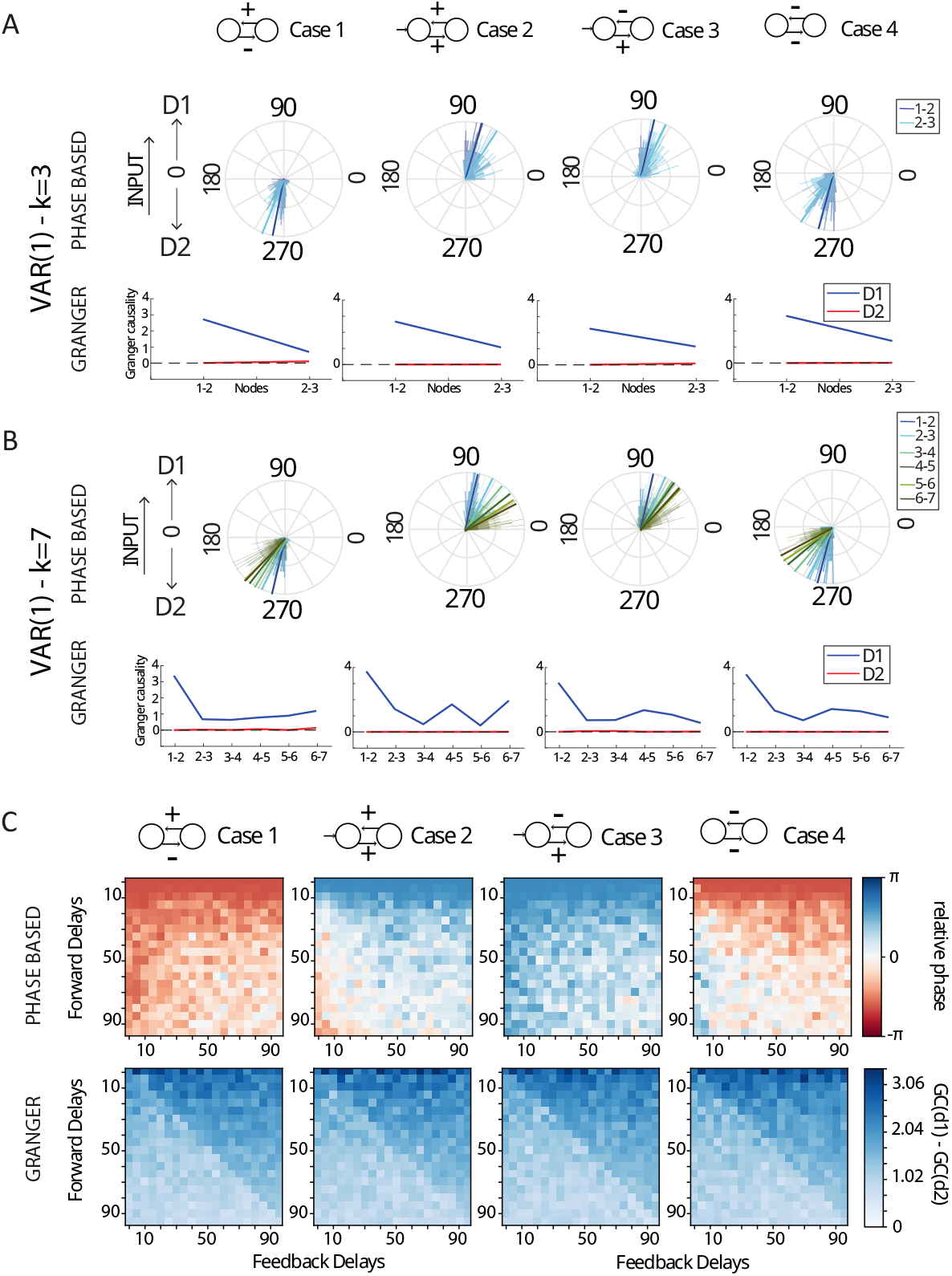
A,B) Results for the VAR(1) case for phase and Granger analysis, for k=3 and k=7 nodes. Each panel shows a case, as described in the upper insert. Each panel is composed of a rose plot illustrating the average phase difference in degrees (90° reveals a flow toward D1, 270° towards D2), and a plot on the right indicating pairwise Granger causality between subsequent nodes. In all cases, Granger values reveal a stronger directionality in the D1 direction, whereas phase differences reveal opposite results than expected in cases 1 and 4. C) Results of the phase-based and Granger analyses for asymmetric delays. Each pixel corresponds to a pair of delays (*δ*_1_, *δ*_2_) for directions D1 and D2. The top row shows the median phase difference across nodes. For visualization purposes, we apply a tanh transformation for the visualization of the relative phases. Case 2 and Case 4 reveal a change of the phase sign when delays of D1 are larger than delays of D2. The bottom row shows the difference between the average of Granger causality coefficients in the two distinct directions (GC(D1)-GC(D2))

### Granger-causality analysis for the 1-D VAR(1) case

We then applied Granger analysis to assess the propagation direction and reveal the causal information flow.

Figure 2A-B shows the GC value in the time domain between all nodes. We found highly similar patterns of GC influences for all four cases, irrespective of the signs of the connections. In particular, GC analyses indicated much stronger Granger-causal influences in Direction 1 (emanating from the perturbed node) as compared to Direction 2, for which Granger-causal influences were negligible. For example, for k=3, we found GC values of 2.566 *±* 0.38 from node 1 to node 2 and 1.019 *±* 0.06 from node 2 to node 3, while GC values in the opposite Direction 2 were smaller than 0.08. Similarly, when k=7, we found the larger GC value from node 1 to node 2 (2.880 *±* 0.31), whereas all other GC values from node *i* to node *i* + 1 (with 2 ≤ *i* ≤ 7) were between 0.925 *±* 0.14 and 0.988 *±* 0.15. By contrast, all the GC values connecting nodes in Direction 2 were smaller than 0.02 *±* 0.02. As expected, most connections did show a significant effect when compared against a theoretical asymptotic null distribution (Barnett and Seth (2014)): This is due to the fact that a small innovation error *ϵ*(*t*) was introduced at all of the nodes.

Together, these results suggest that the Granger and the phase-based analysis provide complementary information: GC reveals the causal flow of information, following the input information propagation starting from the perturbation of Node 1, whereas the second one provides information about the evolution of the phase, which we found to be strongly influenced by the sign of the connectivity.

We generalized these observations to VAR(k) models where *k* ∈ {2, 3, 4} and with different temporal delays (see supplementary materials). Furthermore, we generalized these results to a cortical model grounded in the hierarchical predictive model of inter-areal interactions (Alamia and VanRullen (2019)), which includes biologically plausible temporal delays between subsequent cortical nodes (see supplementary materials). In all cases, GC analyses were consistent with the ground-truth causal information flow, irrespective of the sign of connection weights between nodes.

### Asymmetric Delays

We then investigated, using VAR processes with asymmetric delays, whether the discrepancies between the phase-based and GC analyses are primarily due to the presence of inhibitory connections or can also emerge due to differences in the delays between the two opposite D1 and D2 directions.

For the phase-based analysis, the upper row of Figure 2B shows that results obtained for Case 1 and Case 3 (i.e., opposite signs) are consistent for almost all pairs of delays, with the relative phase representing the averaged phase differences between subsequent nodes. The relative phase change converges to 0 for high temporal delay. However, Cases 2 and 4 (same signs) show an inversion of the phase gradient when the delays in D1 are much greater than the delays in D2. Even though the phase inversion is consistent for multiple pairs of delays, most relative phases for *δ*_1_ *>* 15 and *δ*_2_ *<* 15 are included in a relatively small interval between [−0.08*π*, 0.08*π*] (for clarity, the colorbar is in logscale).

Regarding the Granger analysis (bottom row), all coefficients are positive, confirming an overall flow of information following the direction of the perturbation D1. On the one hand, these results corroborate the conclusion that Granger and the phase-based analysis provide complementary information about the flow directionality, specifically when the connections in the D1 direction are inhibitory. Moreover, they extend these conclusions to the case of asymmetric delays, revealing that very asymmetric delays may further shift the information provided by the two approaches.

### 2D networks and DIFF method

We then analyzed cases in which the nodes are spatially organized in 2D networks and locally connected to their neighbors, as shown in Figure 1B. We considered two distinct conditions in which we continuously perturbed two subsets of nodes: in the first one, the external perturbations affected two non-adjacent nodes (3), in the second one, all the nodes of the lower edge were perturbed (S2). We computed the phase vector field and the Directional Information Flow Field (DIFF) in both conditions for all four cases, and we computed the absolute divergence in both vector fields to identify the source of the external perturbations.

As shown in Figure 3A, the phase analysis provides qualitatively different vector fields in the four cases. In Cases 1 and 3, in which the connections had opposite signs, the phases revealed a flow following the excitatory direction (D2 and D1 in Cases 1 and 3, respectively). In Cases 2 and 4, in which all connections share the same sign, the phases report either an outgoing pattern from the sources in the case of excitatory connections (Case 2) or a more random pattern if all connections are inhibitory (Case 4). To statistically analyze the results, we tested whether the divergence value at the perturbed node was significantly different from that in all other nodes (Figure 3A, lower panels). We consider all divergence values as a univariate Gaussian distribution, and we computed the probability (i.e., the p-value) that the divergence values at the perturbed nodes are sampled from the same distribution. We found significant p-values *p <* 0.05 only for case 2, whereas all other p-values were larger than 0.05. This result was consistent and robust to an increase in the amount of independent Gaussian noise *ϵ* introduced in all nodes, as shown in the upper panels of Figure 3C. This analysis suggests that, in a 2D network, the divergence of the phase-based vector field can reveal the source of propagating activity if all connections are excitatory (Case 2), but not otherwise.

**Figure 3:**
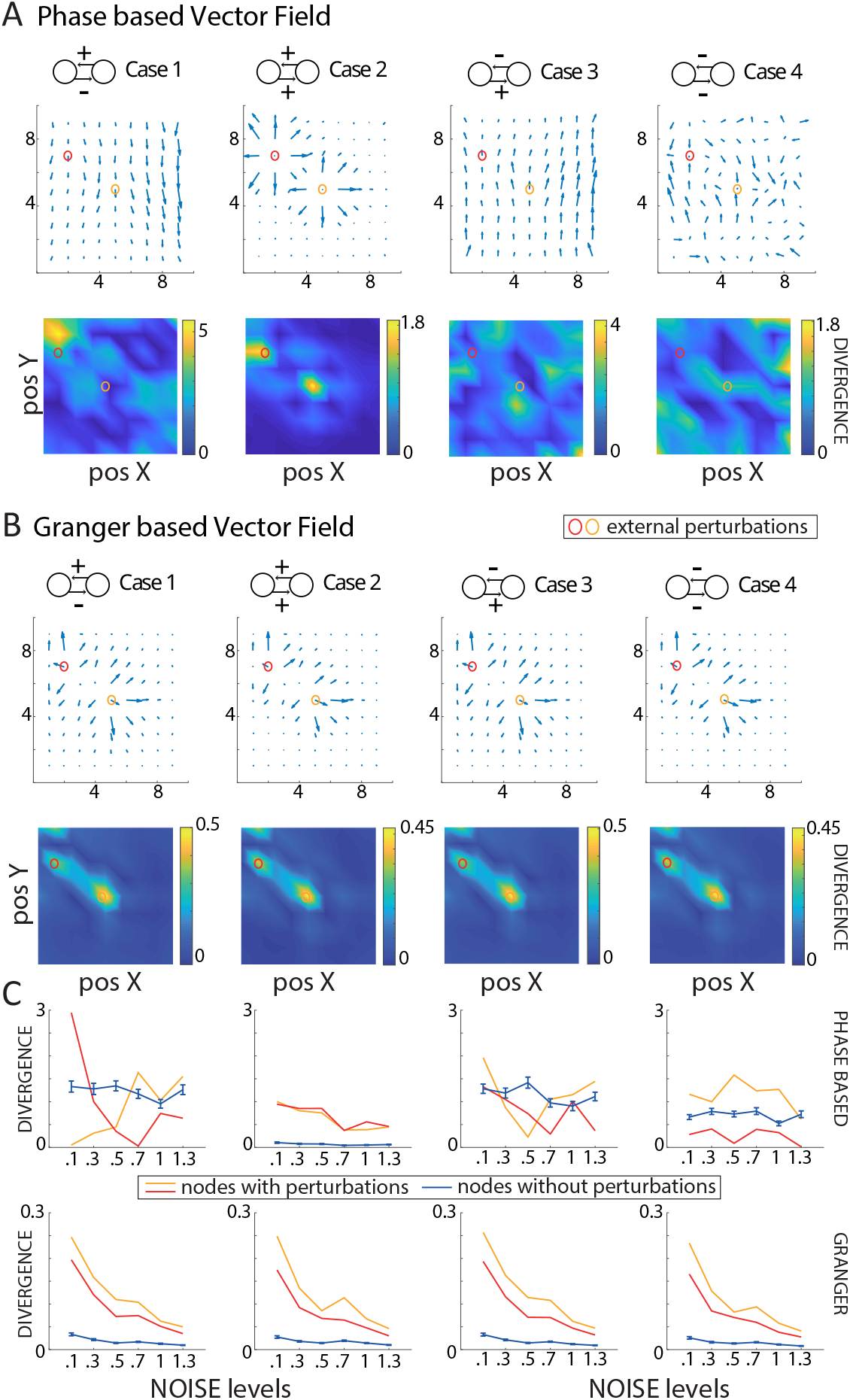
Results for the 2D networks. Panels A and B show the results of the phase-based and Granger-based (i.e., DIFF) vector fields. Each column corresponds to one of the four cases, as represented in each insert above. In each panel, the first and second rows represent the vector field and its divergence, respectively. The two nodes circled in red and yellow are the sources of the perturbations. Panel C) shows the divergence as a function of the noise *ϵ* introduced independently in all nodes. Blue lines are the nodes without perturbation (mean *±* standard deviations), whereas the colored lines represent the divergence in the perturbed nodes.

**Figure 4:**
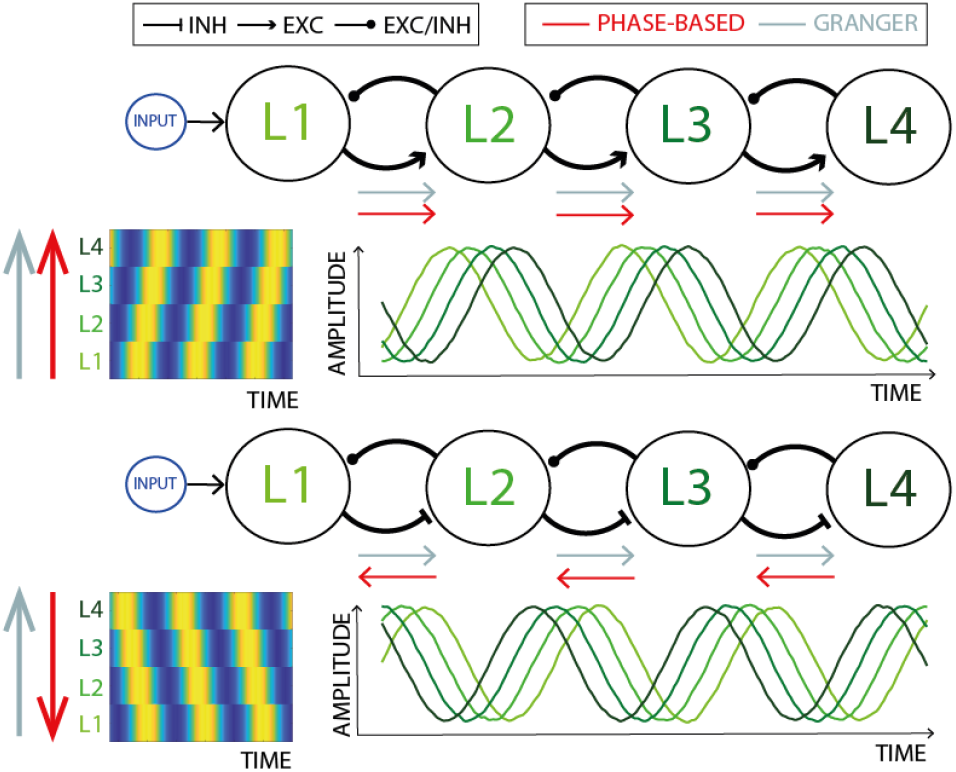
Summary of the results. The upper and lower plots outline the different cases from Figure 1. When the connections of the D1 direction are inhibitory, the phase-based and the Granger-based analyses provide different results.

Contrary to the phase analysis, the Directional Information Flow Field, computed starting from Granger causality measures, provides consistent results across cases (Figure 3B), observing an outward pattern from the external perturbation. In particular, the divergence values at the perturbed nodes are significantly different from the divergence values at all the other nodes for all cases (all *p <* 0.001). Remarkably, these results were consistent with the increase of the noise levels *ϵ* introduced in each node, even though the divergence values at the perturbed nodes decrease with the noise level (Figure 3C), the difference between perturbed nodes and all the other ones remains significant (all *p <* 0.01).

We then confirmed and extended the previous results to the case of a planar wave propagating from the lower edge of the network (akin to Node 1 in the 1D case), as shown in Figure S2. In particular, we observed that the DIFF method applied to the Granger-based vector field reliably identifies the perturbed nodes in all cases, as indicated in Figure S2B, and irrespective of the amount of noise level *ϵ* (S2C). Contrarily, the phase-based vector field reports the correct direction in Cases 2 and 3 (as in the previous 1D analysis), but an opposite pattern in Cases 1 and 4 (specifically closer to perturbed node), similarly to the 1D model. Different from the Granger-based vector field, the divergence of the phase vector field does not identify the source of the planar wave, confirming the complementarity of the two measures.

## Discussion

### Summary of the results

Neural activity manifests itself within and across brain regions as traveling waves Muller et al. (2018); Ermentrout and Kleinfeld (2001); Hatsopoulos et al. (1998); Troy and Shusterman (2007); Zhang et al. (2018) Most methods detect traveling waves by measuring spatial phase gradients, i.e., monotonic and ordered phase changes through space Das et al. (2023); Alexander et al. (2013); Alamia and VanRullen (2019); Muller et al. (2014). The central question of this work was to compare traveling waves, using phase-based method, to the directionality of causal information flow, using Granger Causality. To this end, we analyzed different network configurations of vector autoregressive models, which have been used in several studies for neural simulations and provide a good approximation of neural dynamics (Seth et al. (2015); Vinck et al. (2015); Pourahmadi and Noorbaloochi (2016); Gorrostieta et al. (2013); Ghosh et al. (2019); Marinazzo et al. (2011); He et al. (2019)). Specifically, we investigated different conditions by varying the sign and delay of the connections between nodes. The networks were perturbed by an input to a given node (the first one in the 1D case), causing wave-like patterns across nodes. We demonstrate that the phase ordering in traveling waves does not always correspond to the direction of information flow, as summarized in 4. Specifically, we show that discrepancies can emerge in the case of systems with delayed and/or inhibitory influences. As a methodological solution, we show that Granger causality analysis can, for linear systems, recover the directionality of the information flow. We propose a new measure called DIFF, the Directional Information Flow Field, constructed by analyzing directed causal influences in space and time between neighbours. DIFF yields a vector field on which standard vector calculus measures like Divergence and Curl can be applied. As proof of principle, we show that, in a 2D field where a connected network is perturbed by an external input, the Divergence of DIFF applied to the Granger Causality can identify the spatial source of the perturbations. We will discuss whether causal inference methods provide complementary information to phase-based traveling wave methods in analyzing system dynamics and information flow.

### Biological plausibility of our results

Our results reveal that, under certain conditions, a significant difference can arise in the identified direction of propagation of the activity measured via phase difference and the underlying causal interactions. The results were consistent across different implementations with and without temporal delays and in a model grounded in the Predictive Coding framework (Alamia and VanRullen (2019)). Yet, we note our results were based on linear models with homogeneous connectivity. Considering the first assumption about linearity, one may wonder whether such a discrepancy can be generally found in more biologically plausible systems and whether Granger-causality methods still provide a solution in those cases, given that the cortical dynamics are supposedly generated by non-linear systems (Vinck et al., 2023; Freeman and Skarda, 1985). For non-linear systems, Transfer Entropy methods can be used for the same purpose, and a direct comparison between phase methods and Transfer Entropy needs to be explored in future work.

We further note that linear dynamics may, in many cases, dominate inter-areal connectivity measures due to the strong influence of afferent connections on synaptic potentials in downstream areas (Pesaran et al., 2018; Dowdall et al., 2023; Dowdall and Vinck, 2023; Vinck et al., 2023; Schneider et al., 2021). Studies have also suggested that local oscillatory network dynamics may also be well approximated by linear dynamics, e.g., a damped harmonic oscillatory driven by noise (Spyropoulos et al., 2022).

We argue that the other assumption, namely that the connections between nodes show heterogeneous signs (i.e., both inhibitory and excitatory), may be realistic at a meso- or macro-scopic scale. Regarding cortical networks, while feedforward and feedback projections are glutamatergic (Callaway (2004); Shao and Burkhalter (1996)), we note that feedforward connections target both inhibitory and excitatory neurons, generating both IPSPs and EPSPs in principal cells in a downstream area. However, experimental evidence suggested that feedback activity may have a net suppressive influence on lower areas at a meso/macroscopic level ((Olsen et al., 2012; Isaacson and Scanziani, 2011; Vezoli et al., 2020)). Thus, in our one-dimensional simulation, if the first node represents the activity of a higher area perturbed by an input (e.g., in the case of imagery or a top-down prediction), information would propagate backward from higher regions Rao and Ballard (1999); Rauss et al. (2011). In this case, the discrepancy between Granger-based and phase-based methods would occur if feedback connections have a net suppressive effect (Briggs, 2020; Murphy et al., 1999; Wilmes and Clopath, 2019). In other words, feedback effective inhibition may result in an apparent traveling wave propagating in the opposite, feedforward direction. Furthermore, paradoxical effects of neural activation can arise from balanced E/I networks Tsodyks et al. (1997), as well demonstrated by inhibitory-stabilized networks Ozeki et al. (2009), with long-range excitation leading to decreased activation Chemla et al. (2019b). In particular, high-frequency inputs have been suggested to predominantly target fast-spiking inhibitory interneurons through feedforward connections (Schneider et al. (2023); Spyropoulos et al. (2024); Reynaud et al. (2012)). Interestingly, previous work reported discrepancies between the direction of phase-based and Granger-causal methods in the beta-frequency range (Brovelli et al., 2004a). A possible explanation is that beta is associated with net inhibitory influences (Bastos et al., 2020), causing a discrepancy between phase-based on Granger-causality methods.

In light of these considerations, our conclusions call for caution when interpreting experimental results about the propagation of information in neural recordings, which may lead to wrong assumptions about the effective flow of information.

### Complementary of Granger Causality and phase-based analyses

Granger Causality analysis has been widely used in the neurosciences for many applications (Marinazzo et al. (2011); Brovelli et al. (2015, 2004b); Marinazzo et al. (2008); Shojaie and Fox (2022); Cekic et al. (2018); Stokes and Purdon (2017); Seth et al. (2015); Ding et al. (2006)); however, Granger-causality has not been applied to analyze complex spatiotemporal patterns. When applied to a linear system, Granger-causality provides a well-defined solution with uniquely separated contributions (Seth et al. (2015); Marinazzo et al. (2008)), and reliably captures the causal directionality following the propagation of the input provided to the system. We argue that Granger-causality can capture the ground truth of causal influences for a linear system of coupled stochastic processes, in which the causal influences across nodes can be uniquely decomposed, and where the concept of causal influence has a well-established mathematical definition (**?**). Evidently, the phase pattern provides complementary information as an emergent phenomenon that is functionally relevant. For example, the phase pattern might encode information that can be extracted in single trials (Muller et al., 2014; Rubino et al., 2006), and may further sculpt the interactions or plasticity between nodes, and predict behavior (Benigno et al., 2023; Davis et al., 2020). The phase can also provide information about the speed of a wave traveling in a given direction. It is also important to emphasize that Granger-based methods have several limitations in general. First, uncorrelated or correlated noise can strongly affect Granger-causality inferences. For this problem, several solutions have been proposed, e.g., time-reversal controls (Vinck et al., 2015; Haufe et al., 2014) or Kalman-filtering approaches (Nalatore et al., 2007). Second, Granger-causality is explicitly defined for stochastic time series, and can therefore not be used to infer causality for deterministic systems. Topological methods may, in such cases, allow for a definition and measurement of causality (Sugihara et al., 2012). Third, if a system is strongly nonlinear, then nonlinear methods like Transfer Entropy are required to capture causal relationships (**?**). In this case, the same framework and methodology can be applied using Transfer Entropy. Alternatively, more direct modelling approaches to capture the underlying dynamical system, e.g., dynamic causal modelling, could be suited for such cases. More generally, macroscopic traveling waves can emerge from an intricate propagation of activity in a complex underlying network (Budzinski et al., 2023), whose mechanistic origin may be hard to infer. That is, the mechanistic origin of phase gradients at the macroscopic level may have a much more complex origin than a simple propagation of a single perturbation, e.g., axonal propagation. Fourth, given the presence of widespread connections in cortex (Vezoli et al., 2020), identifying causal relationships may require a multivariate approach. Fifth, Granger-causality is not designed to be applied to a unique trial but rather to a set of many single trials, as it relies on fitting an underlying generative model. By contrast, phase-based methods can be applied to unique trials. It is possible, however, that there are distinct patterns of traveling waves that can be detected in different sets of trials, which can then be subsequently analyzed separately with Granger-causality analyses. In this way, phase-based and Granger-causality measures can be applied synergistically.

Discrepancies between Granger-causality and phase-based analysis may also emerge because of the unique properties of electrophysiological data. Importantly, the sign of LFP/EEG electrophysiological data is arbitrary and depends on the reference, and is subject to dipole reversals (Pesaran et al., 2018). Hence, the phase of electrophysiological signals depends on referencing schemes and the underlying generative model of the electric field. Granger-causality, on the other hand, is invariant to a scaling of signals and sign reversal. Thus, the DIFF analysis can, in principle, also be applied to current-source-density data or bipolarly derived data.

Future work will be needed to explore more realistic modeling, which would allow us to better capture the richer cortical dynamics while also fully exploring the range of parameter spaces, including the strength of the connection and different temporal delays. This more comprehensive approach would allow us to take one step closer to proper inference and provide a complete outlook on the methods to quantify waves’ propagation.

## Methods

### General VAR models

Consider a *k*-variate VAR model of order *T*, described by the system of discrete-time equations:

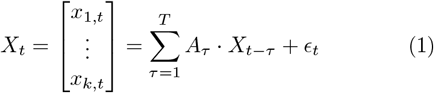

Here, *X*_*t*_ is a vector describing the evolution of a set of k variables. *A*_*τ*_ represents a (*k×k*)-coefficient matrix that describes the influence of the past state at lag *τ* (*X*_*t*−*τ*_) on the current state. The innovation term *ϵ*_*t*_ is a vector of the same dimension as *X*_*t*_ where each component is assumed to be white noise with zero mean and covariance matrix Σ.

Next, consider a simplification of this system, in which the nodes are sequentially organized, such that there are only direct interactions from a node *i* with the nodes *i* − 1 and *i*+1. We call the flow from node *i* to node *i*+1 “Direction 1” (D1), which is the information flow caused by the original perturbation. For a biological system, this may be either the forward connection (e.g., a sensory stimulus) or a feedback connection (e.g., imagery). We call the flow from node *i* + 1 to node *i* “Direction 2” (D2). In the system analyzed, *∀ τ* ∈ [1, *T*], *A*_*τ*_ is a tridiagonal matrix with a specific set of coefficients defined as:

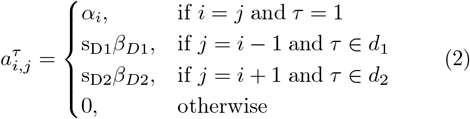

where *α*_*i*_, *β*_*D*1_, *β*_*D*2_ ∈ (ℝ^+^)^3^ are random variables drawn from 𝒩_trunc+_(0, 2), a Gaussian distribution truncated for all values below zero, s_D1_, s_D2_ ∈ {−1, 1}^2^ determine the sign of the two opposite directions. and *d*_1_ and *d*_2_ are sets of delays that define connectivity at specific lags. Note that there is a unique weight, *β*_1_, connecting neighboring nodes in direction D1. Similar connectivity applies for D2, but the values of *α*_*i*_ depend on the nodes.

If we take a 2-variate VAR model, this yields in total 4 possible cases illustrated in Figure 1A, conditional on the value of s_D1_ and s_D2_. In the following, we will first consider the simplest VAR(1) case. Then, we will introduce models with higher orders, i.e., from VAR(2) to VAR(4), and with asymmetric delays. After a section describing a biologically plausible cortical model, we extend our results from 1D to 2D cases.

### The VAR(1) case

Starting from the general case described above, we first consider the condition with *d*_1_ = *d*_2_ = 1, i.e., using only the previous state to describe the current one :

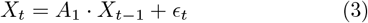

In this case, we test models with k=3 or k=7 nodes. For each of the four cases described above (Figure 1A), we run 1000 simulations of 500 time steps each. In each simulation, we determine the stability of the model by assessing that all eigenvalue modules associated with the companion matrix were |*λ*_*i*_ |*<* 1. We discard all runs in which the system was unstable.

The first node is fed an input signal, a series of independent, pseudo-random values drawn from a Gaussian distribution 𝒩 (0, 1). In all nodes, we introduced independent Gaussian noise *ϵ*(*t*) from 𝒩 (0, 0.1).

For all analyses, we only considered stable models. We first assessed the stability of the 1000 randomly generated models for each of the four conditions. We found that around 35% of simulations were stable for *k* = 3 and 25% for *k* = 7 for the four conditions.

### From VAR(1) to VAR(4)

We also consider the case with multiple temporal delays in a VAR model with order *p >* 1 (as in Equation 1).

Specifically, we investigated different conditions with specific sets of symmetric delays (i.e., *d* = *d*_1_ = *d*_2_) for information transfer between nodes. We study 3 different VAR models with *d* = {1, *δ* }, where *δ*∈ { 2, 3, 4} . In addition, we also considered the case where *d* = {1, 2, 3, 4} . The order of these VAR processes is given by max(*d*)

All the other parameters (i.e., number of simulations, time-steps, nodes, noise, and stability assessment) were the same as in the case without temporal delays. For all simulations, we assessed the stability of the 1000 simulations we performed with randomized coefficients. Overall, around 20% and 15% of the simulations were stable for *k* = 3 and *k* = 7, respectively. When considering several delays at once (*d* = {1, 2, 3, 4}), the stability dropped to less than 10% of the simulations, leaving, however, more than 50 runs in the lowest cases (cases 2 and 4). As previously, we considered only stable models for the analyses.

### Asymmetric Delays

We also study models with asymmetric delays (i.e., *d*_1_ = {*δ*_1_}, *d*_2_ = {*δ*_2_} and *δ*_1_ ≠ *δ*_2_). We simulate all combinations of models with delays varying from 1 to 100. Because of long delays, we want to run the phase-based and the Granger analysis once all nodes have been perturbed by the input, i.e., after time *t* = (*d*_1_ + *d*_2_) * *k, k* being the number of nodes. For this reason, we increased the simulation length from 500 to 2000 time steps- and keep only the relevant part of the simulation. The results regarding both the phase-based and the Granger-based analysis were computed by taking the median of the phase/Granger values differences across all nodes for 50 stable simulations in each case.

### Simulations for a predictive coding model

We also apply our analysis to a biologically plausible model. Based on the generic VAR model given in Equation 1, we can introduce a specific set of parameters to match a given connectivity matrix or other existing models. Here, we set the parameters to match the dynamical system described in Alamia and VanRullen (2019).

We started from the equations of a two-layer system as described in (Alamia and VanRullen, 2019):

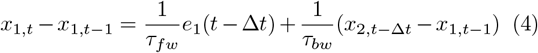

where *τ*_*fw*_ and *τ*_*bw*_ are time constants, *e*_1_(*t*) is the prediction error in the first layer:

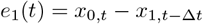

and *x*_0,*t*_ is the bottom-up input to the network at time *t*. This, eventually, simplifies to:

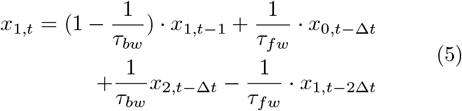

For this dynamical system, we have an equivalent VAR(2Δ*t*) model where all coefficient matrices have zero values except for *A*^1^, *A*^Δ*t*^, and *A*^2Δ*t*^. Thus, we have:

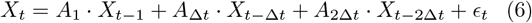

To match the dynamical system given by Equation 4, we need *A*_1_ to be a diagonal matrix with the diagonal coefficients equal to 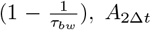 to be another diagonal matrix with the diagonal coefficients equal to 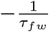 and *A*_Δ*t*_ to be a tridiagonal matrix with 0 on the diagonal, 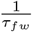 on the lower diagonal and 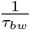 on the upper diagonal. As in Alamia and VanRullen (2019), we set *τ*_*fw*_ = 30 ms and *τ*_*bw*_ = 200 ms, these values granted a stable behavior and can be interpreted from a biological perspective as time constants. From this VAR(2Δ*t*) model, one can compute the frequency and oscillatory behavior, which depend on its parameters *τ*_*fw*_, *τ*_*bw*_ and Δ*t*, as demonstrated in Alamia and VanRullen (2019). Note that Equation 6 describes the 2-layer model given by Equation 5 but does not hold any assumption on the dimensionality of X. It can be easily generalized to *k* nodes by increasing the size of vector *X* and of the coefficient matrices *A* with the same structure as the one described in this case.

Considering this implementation, we tested models having *k* = 4 nodes. For each of the four cases described above (Figure 1A), we run 300 simulations of 500-time steps with independent Gaussian noise *ϵ*(*t*).

### Quantifying waves’ direction of propagation

In all simulations in which the model was stable (i.e., all eigenvalues |*λ*_*i*_ | *<* 1), we used two different approaches to quantify the activity’s direction of propagation. The first method was based on the estimates of the signal’s instantaneous phase; the second one applied Granger Causality to assess the flow’s direction.

### Phase-based analysis

At the end of each simulation, we computed the Hilbert transform *H*(*x*_*k*_) of each signal *x*_*k*_ for each *k* node and calculated the complex vector obtained by computing the phase difference between subsequent nodes at any time point *t*:

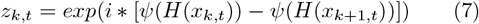

where *ψ*(*H*(*x*_*k,t*_)) is the phase of the Hilbert-transformed signal *x*_*t*_ at the node *k*. We then sum all complex vectors over time and compute the average phase difference between nodes:

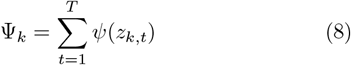

The sign of Ψ_*k*_ provides the phase shift between subsequent nodes: if positive, it indicates a wave propagating from node *k* to node *k* + 1; vice versa, a negative value reveals activity propagating from the higher to the lower nodes.

### Granger analysis

We used the multivariate Granger causality analysis (MVGC) toolbox in MATLAB (MathWorks) to compute the Granger causality between subsequent nodes in all cases (Barnett and Seth (2014)). For each condition and stable model, we first fit the model’s parameters, considering each simulation as a trial. We estimated the best model’s order by optimizing the Akaike Information Criterion (we obtained similar results using the Bayesian Information Criterion). For the 1-D VAR(1) case, we fitted the model’s order for k=3 and k=7 nodes, optimizing Akaike’s criterion, and we obtained a fit of the third and fifth order, respectively, for all conditions. The results reported did not change qualitatively by imposing the model’s order to equal 1 before fitting the model’s parameters.

Besides estimating the model’s parameters, such a fit provides the covariance matrix of the residuals, which is next used to calculate the autocovariance sequence according to the estimated VAR model. Lastly, the pair-wise time-domain Granger causalities are computed for each pair of *x*_*i*_ from the covariance sequence. Significant effects were estimated against the null hypothesis of zero causality based on a theoretical asymptotic null distribution (Barnett and Seth (2014)).

### Generalization to 2D networks

#### Nodes configuration and simulations

We generalized our results from the one-dimensional case, i.e., a series of consecutively connected nodes, to the two-dimensional case, in which the nodes are organized in a square grid and connected to their neighbors. We considered squared grids of side *k × k*, with *k* = 9. It is possible to formalize the 2D grid arrangement as a 1D case with long-range connections (i.e., between non-subsequent nodes). As specified in Figure 1C, the parameters connecting node *i* to node *i* − *k* determine D1, whereas parameters from node *i* to node *i* + *k* determine D2. The parameters between node *i* and its neighbors *i* ∓ 1 characterize the lateral connections. As in the 1D case, we can fully describe the model via the parameters of the matrix *A*_*τ*_ . For this 2D case, we focus on a VAR(1) process, and we can describe *A*_1_ as:

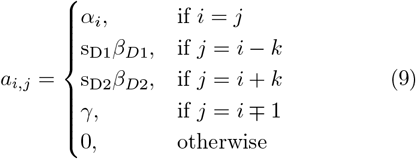

where *α, β*_1_, *β*_2_, *γ* ∈ (ℝ^+^)^4^ are random variables drawn from 𝒩_trunc+_(0, 2), a Gaussian distribution truncated for all values below zero, and s_D1_, s_D2_ ∈ {−1, 1 }^2^ determine the sign of the two opposite directions. Considering this configuration, we tested 300 simulations of 100 time-steps in which we perturbed a subset of specific nodes either at different locations, or at the lower edge of the grid, with a series of independent, pseudo-random values drawn from a Gaussian distribution 𝒩(0, 2). In all cases, we introduced independent Gaussian noise *ϵ*(*t*) from 𝒩(0, *η*) at each node. The value of *η* was modulated to modulate the noise levels over different simulations.

#### Phase-based analysis

We performed the phase-based analysis by estimating a phase-based vector field in each grid. In particular, for all nodes *k* we computed the phase differences between the Hilbert transforms of each signal *H*(*x*_*k*_) and the neighboring nodes whose distance to node *k* was smaller than or equal to 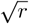. To consider the circle capturing direct diagonal neighbors as well as directly connected neighbors, we set *r* = 2. Such a phase difference was defined as:

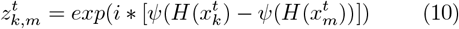

where 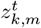 is the phase difference between the *k* node and its neighbor *m* at time *t*. For all neighbors *m*, we computed the sinusoid of such phase difference averaged across time points:

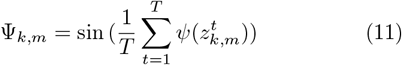

Lastly, we average the phase difference Ψ_*k,m*_ over all neighbors *m*, considering their relative position with respect to the node *k*. This results in a vector field in which each vector, centered at every node, points to the mean phase difference between its neighbors whose distance to *k* is smaller than or equal to 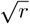.

#### DIFF method

Similar to the phase-based vector field, we computed a vector field based on the directional information flow estimated via Granger measures. The Directional Information Flow Field (DIFF) is then a vector field obtained by computing the pair-wise Granger causality between every node and its neighbors whose distance to *k* was smaller than or equal to 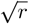 (as before, we set *r* = 2).

For every node *k*, we averaged the Granger values over all neighbors *m*, considering their relative position with respect to the node *k*. This generates a vector field *GC*_*x,y*_ where, at each node’s position *x, y*, the vector points in the averaged information flow estimated as the mean Granger value between its neighbors. As above, all Granger analyses were computed using the MVGC toolbox in Matlab (Barnett and Seth, 2014). Finally, for both the phase-based and the Granger-based vector fields *V F*_*x,y*_, we estimated the absolute divergence at every node as:

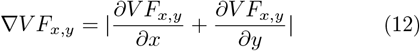

Crucially, values of *V F*_*x,y*_ larger than 0 locally identify the source of the external perturbations.

## Acknowledgements

The authors wish to thank Lyle Muller and Laura Dugué for their helpful and constructive comments. AA, FC and MV conceived of the study. AA and AG analyzed data. AA and MV wrote the paper, with comments from the other authors. MV was supported by an ERC starting grant (850861) SPATEMP; DFG VI Grants (908/5-1 and 908/7-1); an NWO VIDI Grant; and the Dutch Brain Interface Initiative (DBI2). AA was supported by ERC starting grant (OSCI-PRED, no. 101075930). Views and opinions expressed are, however, those of the authors only and do not necessarily reflect those of the European Union or the European Research Council (ERC). Neither the European Union nor the granting authority can be held responsible for them.

## Supplementary: All simulations with 1D networks

Below, we describe the results for all the simulations in the one-dimensional case. We generalized and replicated our results to VAR(k) models, where *k* ∈ {2, 3, 4}, and to a cortical model grounded in the hierarchical predictive model of inter-areal interactions (Alamia and VanRullen (2019)), which includes biologically plausible temporal delays between subsequent cortical nodes.

## Supplementary Materials

### Supplementary Figure 01: All simulations with 1D networks

#### From VAR(1) to VAR(4)

We tested whether the effect could be generalized to cases with different sets of temporal delays. First, we assessed the stability of the 1000 simulations we performed with randomized coefficients. Overall, around 20% and 15% of the simulations were stable for *k* = 3 and *k* = 7, respectively. When considering several delays at once (*d* = { 1, 2, 3, 4 }), the stability dropped to less than 10% of the simulations, leaving, however, more than 50 runs in the lowest cases (cases 2 and 4). As previously, we considered only stable models for the subsequent analyses.

#### Phase-based analysis

As in the case without temporal delays, we found consistent results for k=3 and k=7 nodes (see Figure S1A,B for the VAR(2) case). In particular, we found a backward propagation in the first and fourth cases, as confirmed by a negative phase difference between all subsequent nodes, but the opposite direction in the second and third cases, corroborating our previous results. These results were very consistent across the different delays. As in the VAR(1) case, a Von Mises test (V-test) for non-uniformity of circular data considering the mean direction in any pair of nodes confirmed the results: in all conditions and pair of nodes, we obtained for k=3 all V-value *>* 120 and *p <* 0.0001 when considering *τ* = [1, 4], and V-value *>* 65 and *p <* 0.0001 when having several delays. Similarly, for k=7 and *τ* = [1, 4], we obtained all V-value *>* 50 (with larger values *≈* 110 for lower nodes, but invariably significant) and *p <* 0.0001, and V-value *>* 20 when considering several delays.

As in the case without temporal delays, the results of our simulations corroborated the finding that a positive sign from the node *i* to node *i* + 1 produces a phase difference consistent with D1 direction (i.e., a forward flow from the input), whereas a negative one leads to a phase difference corresponding to a flow in D2 direction (towards the input).

#### Granger analysis

The Granger analysis was applied to all nodes’ activity to estimate the causal propagation direction. We fitted the model’s order for k=3 and k=7 nodes for each temporal delay, optimizing Akaike’s criterion. For a single temporal delay, i.e., *τ* ∈ {2, 3, 4 }, we obtained orders between 3 and 5, and we obtained an order equal to 8 when considering all the temporal delays, i.e., *τ* = {1, .., 4}.

Considering the k=3 case, we found very similar time-domain Granger Causality (GC) values when having a single temporal delay. For instance, when *τ* = 2, we found 1.377 *±* 0.10 from node 1 to node 2, and 0.88 *±* 0.20 from node 2 to node 3, and 0.284 *±* 0.04 from node 1 to node 3. All other GC values were smaller than 0.05. When considering the case with all temporal delays, we obtained relatively smaller values: 0.804 *±* 0.04 from node 1 to node 2, and 0.56 *±* 0.05 from node 2 to node 3, and 0.10 *±* 0.02 from node 1 to node 3.

Regarding k=7 and *τ* = 2 (very similar values were observed in the other cases having one single temporal delay), we found the larger GC values in lower nodes: progressively from node 1 to node 7 we reported 1.433 *±* 0.31, 0.768 *±* 0.11, 0.16 *±* 0.06, and then all other values were smaller than 0.07 *±* 0.03. Similarly, when *τ* = {1, .., 4 }, we found value decreasing from lower to higher nodes: 0.876 *±* 0.08, 0.346 *±* 0.03, 0.097 *±* 0.2, and then all other values were smaller than 0.05 *±* 0.02.

Altogether, the Granger analysis confirms that the causal flow of information propagates the input information from the perturbed node 1 to other nodes (Direction D1), irrespective of the sign of the *a*_*i,i*+1_ coefficient.

**Figure S1:**
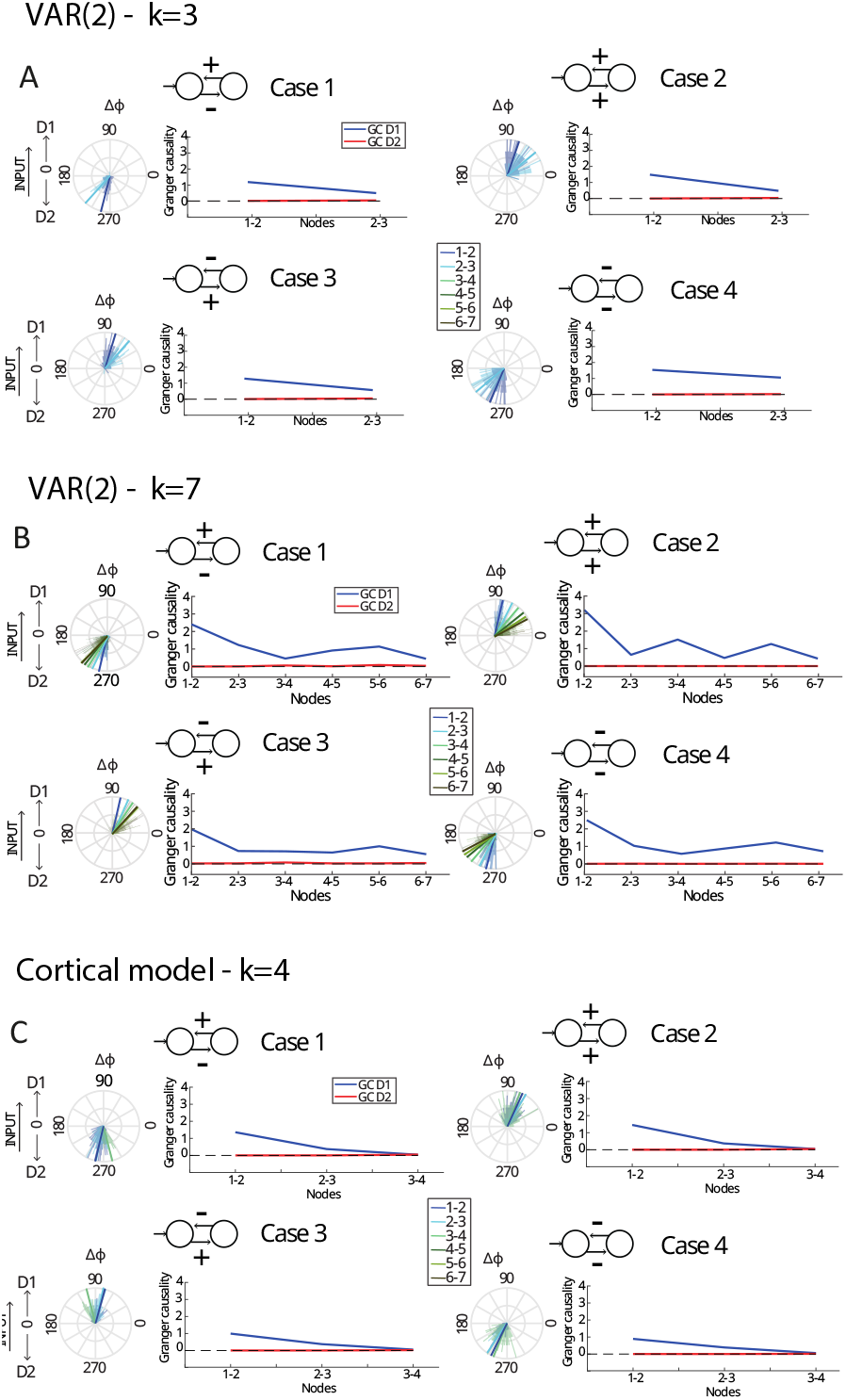
Results for the VAR(2) condition with k=3 nodes (panel A) and k=7 nodes (panel B). Panel C shows the results for the cortical relevant model (based on predictive coding dynamics). Each panel shows the four cases, as specified in the upper insert. For each case, a rose plot illustrates the average phase difference in degrees (90° reveals a flow toward D1, 270° towards D2), and the plot on the right indicates pairwise Granger causality between subsequent nodes. In all panels and simulations, the results show that Granger values consistently reveal the expected directionality (i.e., D1), whereas phase differences consistently reveal opposite results in cases 1 and 4.

#### Simulations for a predictive coding model

Next, we generalized our results to a model grounded in the predictive coding framework (Alamia and VanRullen (2019)), which includes biologically plausible temporal delays between subsequent nodes. We simulated 300 instances of the model for 500 time steps for each of the four conditions. The model had *k* = 4 nodes, and its parameters were chosen to be biologically interpretable and with a stable behavior (see Methods, and Alamia and VanRullen (2019); Schwenk and Alamia (2024)).

#### Phase-based analysis

The results based on the phase analysis support the same conclusions as in the previous simulations, as shown in Figure S1C. In particular, we observed opposite propagation directions depending on the sign of the *a*_*i,i*+1_ coefficient. In the first and fourth cases, when the *a*_*i,i*+1_ coefficient is negative, we observe a consistent, negative phase difference across subsequent nodes, revealing a propagation along direction D2. On the other hand, a positive *a*_*i,i*+1_ coefficient (i.e., second and third cases) generates the opposite direction D1, as confirmed by a positive phase difference between subsequent nodes. These results were confirmed by a V-test for non-uniformity of circular data performed over the 300 simulations and considering the mean direction in any pair of nodes (in all conditions and pair of nodes V-value*>* 280 and *p <* 0.0001).

#### Granger analysis

We then estimated the Granger causalities between all subsequent nodes. We assessed the best model’s order for each condition, having an order of 19 for the first condition and 18 for all the other ones. Figure S1C illustrates the time-domain Granger Causality (GC) values between all nodes. Asterisks reveal a significant effect when compared against a theoretical asymptotic null distribution (Barnett and Seth (2014)). Across conditions, we found decreasing GC values across the hierarchy: 1.407 *±* 0.36 from node 1 to node 2, 0.340 *±* 0.03 from node 2 to node 3, and 0.05 *±* 0.01 from node 3 to node 4. All other GC values (i.e., in the opposite direction and between non-subsequent nodes) were smaller than 0.02 *±* 0.01.

## Supplementary simulations with Planar 2D traveling waves

As reported in the main text, we also investigate the case of a planar wave propagating from the lower edge of the network (akin to node 1 in the 1D case), as shown in Figure S2. Similarly to the case with localized sources generating radial waves, also a planar wave can be reliably identified by the DIFF method applied to the Granger-based vector field, irrespective of the amount of noise level *ϵ* (S2C). Different from the Granger-based vector field, the divergence of the phase vector field does not identify the source of the planar wave in all cases.

**Figure S2:**
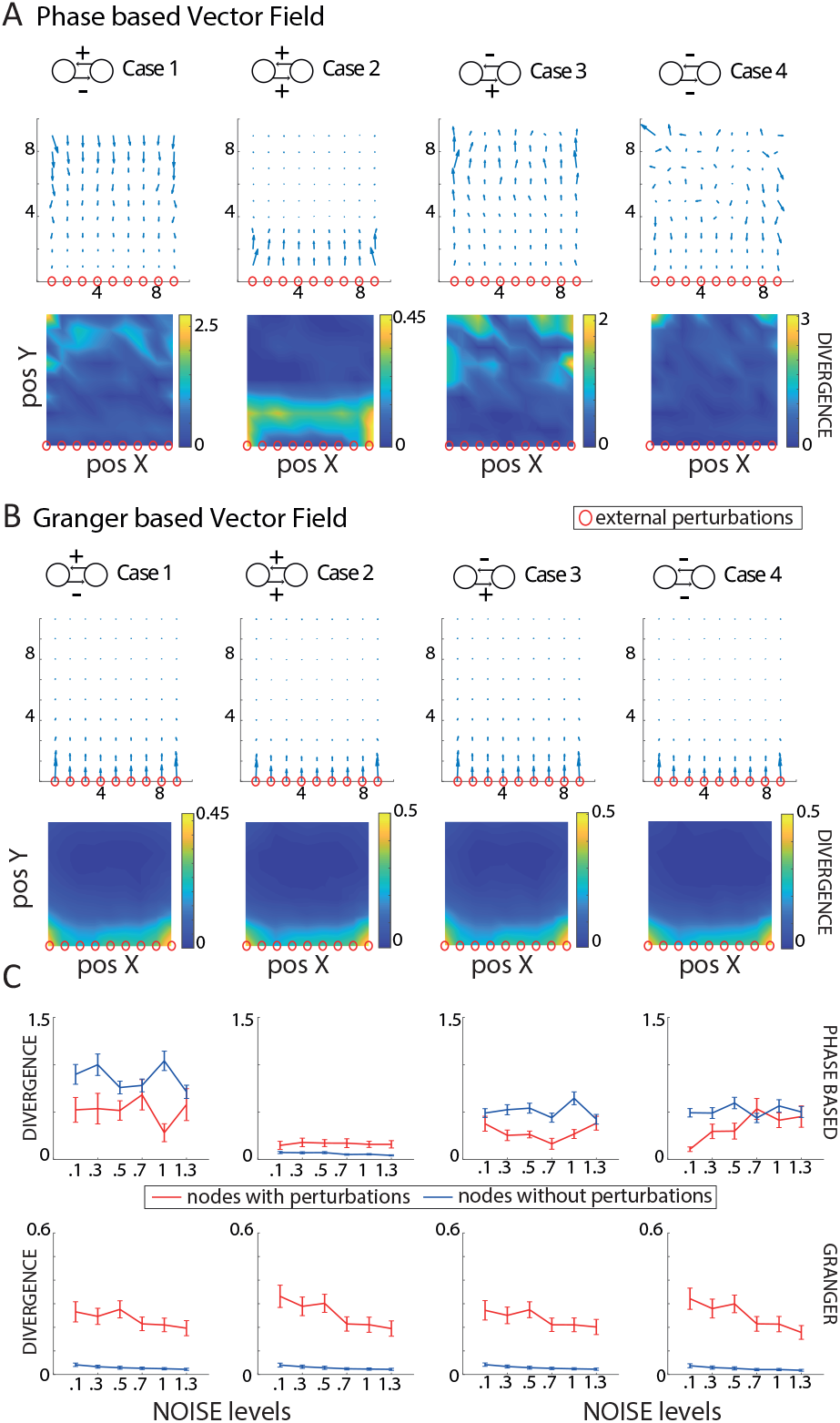
Results for the 2D networks when perturbing the lower edge of the network. Panels A show the results of the phase-based vector field for the four cases, as summarized above each column. The first line represents the vector field, and the second one its divergence. The red nodes are the sources of the external perturbations. Panel B shows the results for the Granger-based (i.e., DIFF) vector fields, same organization as in panel A. Panel C) shows the changes in the divergence as a function of the noise *ϵ* introduced independently in all nodes. Blue lines are the nodes without perturbation (mean *±* standard deviations), whereas the red lines represent the divergence in the perturbed nodes.

